# MARK2 phosphorylates KIF13A at a 14-3-3 binding site to polarize vesicular transport of transferrin receptor within dendrites

**DOI:** 10.1101/2023.07.11.548513

**Authors:** Yue Han, Min Li, Bingqing Zhao, Huichao Wang, Yan Liu, Zhijun Liu, Jiaxi Xu, Rui Yang

**Affiliations:** Institute of Neuroscience, Translational Medicine Institute, Health Science Center, Xi’an Jiaotong University, 710061 Xi’an, China; Department of Physiology and Pathophysiology, School of Basic Medical Sciences, Health Science Center, Xi’an Jiaotong University, 710061 Xi’an, China; Shaanxi Provincial Center for Regenerative Medicine and Surgical Engineering, First Affiliated Hospital of Xi’an Jiaotong University, 710061 Xi’an, China; The Jungers Center for Neurosciences Research, Oregon Health & Science University, Portland, Oregon

**Keywords:** Neuron polarity, Selective transport, TfR vesicle, Kinesin-3 family, KIF13A, MARK2

## Abstract

Neurons regulate the microtubule-based transport of certain vesicles selectively into axons or dendrites to ensure proper polarization of function. The mechanism of this polarized vesicle transport is still not fully elucidated, though it is known to involve kinesins, which drive anterograde transport on microtubules. Here we explore how the kinesin-3 family member KIF13A is regulated such that vesicles containing transferrin receptor (TfR) travel only to dendrites. In experiments involving live-cell imaging, knockout of KIF13A, BioID assay, we found that the kinase MARK2 phosphorylates KIF13A at a 14-3-3 binding motif, strengthening interaction of KIF13A with 14-3-3 such that it dissociates from TfR-containing vesicles, which therefore cannot enter axons. Overexpression of KIF13A or knockout of MARK2 leads to axonal transport of TfR-containing vesicles. These results suggest a novel kinesin-based mechanism for polarized transport of vesicles to dendrites.

**Significance:** Our findings suggest that at least one type of vesicles, those containing transferrin receptor, travel exclusively to dendrites and are excluded from axons because the kinase MARK2 phosphorylates the kinesin KIF13A to promote its separation from vesicles at the proximal axon, preventing vesicle transport into axons, such that they travel only to dendrites. Future studies should explore how this mechanism of polarized vesicle transport supports neuronal function.

## Introduction

Neurons are polarized into axonal and soma-dendritic domains to ensure unidirectional information processing. This polarization arises, in part, through the selective trafficking of vesicles along microtubules into one domain or the other: vesicles containing dendritic proteins are transported efficiently into dendrites but do not enter the axon, while vesicles containing axonal proteins preferentially travel to axon even if they are not actively excluded from dendrites (1, 2). Kinesins, which act as motors to drive anterograde transport along microtubules, appear to help mediate polarized vesicle transport in neurons (3–6), but the details of how they accomplish this remain unclear.

Neurons express at least 15 kinesins, and different kinesins appear to help transport different types of vesicles. We previously showed while vesicles transported by the kinesin-1 family member KIF5A or KIF5C preferentially enter axons, the kinesin-3 family member: KIF13A transported vesicles move largely within dendrites, but not axons. KIF13A transported vesicles containing mannose-6-phosphate receptor (7), viral proteins, serotonin type 1A receptor (8) or transferrin receptor (1, 9). KIF13A also helps the α-amino-3-hydroxy-5-methyl-4-isoxazole-propionic acid (AMPA) receptor translocate to dendritic spines during long-term potentiation (10). Given the observations that KIF13A drives anterograde transport along microtubules and that axons contain abundant microtubules initiated from the cell body with their “+” ends pointing to the distal axon, we wondered why KIF13A-transported vesicles do not enter axons.

Here we examined this question by knocking out or overexpressing KIF13A in primary neuronal cultures and examining the effects on transport of vesicles containing transferrin receptor (TfR). TfR is a ubiquitous transmembrane glycoprotein that mediates iron uptake from circulating transferrin (Tf) at the plasma membrane. The TfR-Tf complex formed at the cell surface is then delivered to endosomes (11). In neurons, vesicles containing TfR are mainly formed and transported at the soma-dendritic area (12). Therefore, it is often used to study dendritic selective transport. In our study, we found that KIF13A is a major transporter for vesicles containing TfR at the soma-dendritic area. Microtubule affinity-regulating kinase 2 (MARK2/Par1) phosphorylation of KIF13A-tail at the 14-3-3 binding motif is required for keeping TfR vesicles dendritically. Overexpression of KIF13A or knockout of MARK2 leads to axonal transport of TfR-containing vesicles.

## Results

### KIF13A transports vesicles containing transferrin receptor to dendrites

As a confirmation of our previous work (1), GFP-fused KIF13A-tail with Halo-tagged TfR were co-expressed in cultured hippocampal neurons and the movement of vesicles were recorded with time-lapse spinning disc microscopy (**Figure 1A-B, Movie 1**). Vesicles containing KIF13A-tail and vesicles containing TfR were overlapped and were bi-directionally moving in dendrites. Very few vesicles were observed entering axons.

**Figure 1.**
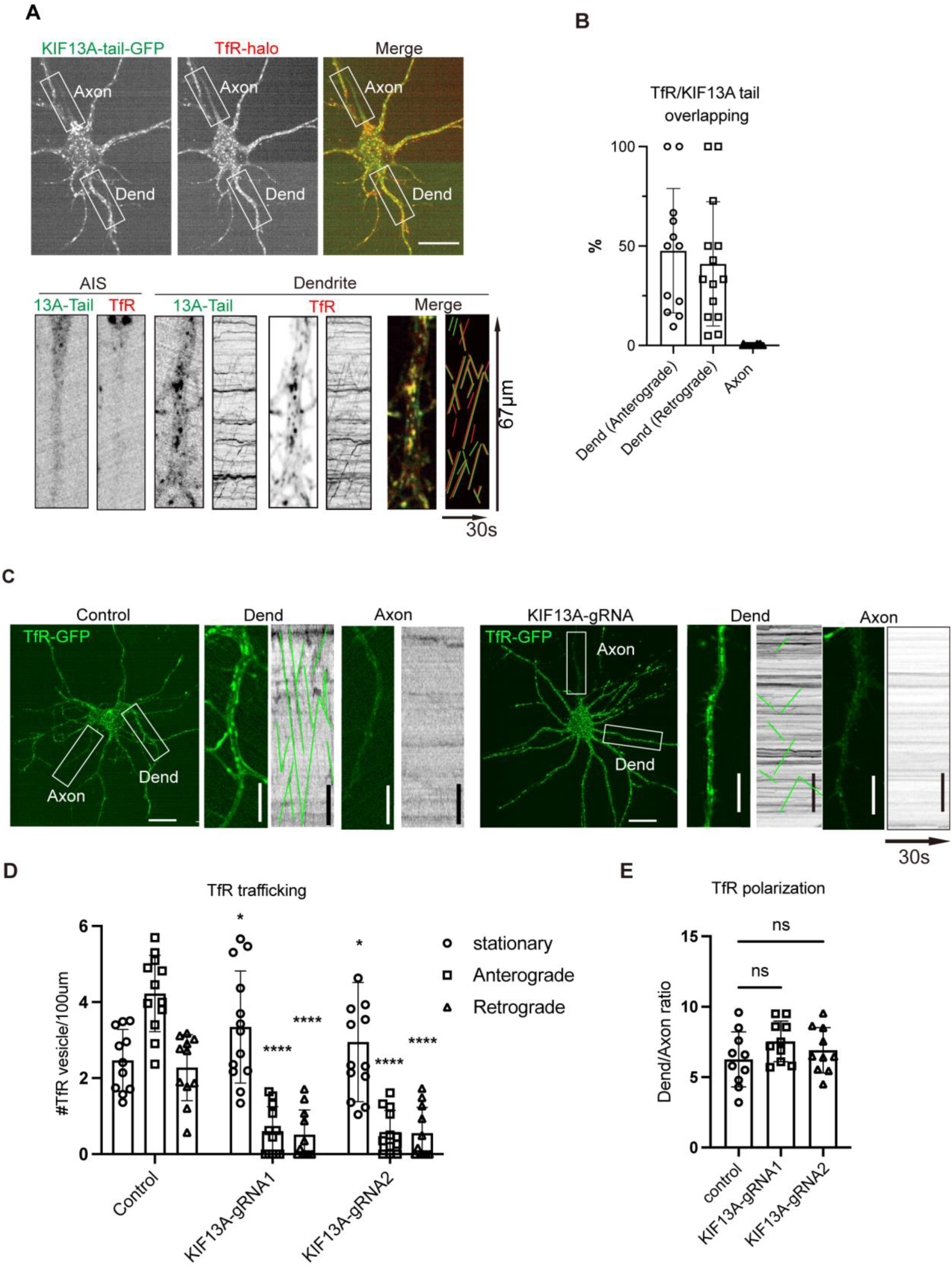
KIF13A drives polarized transport of vesicles containing transferrin receptor (TfR) into dendrites. **(A-B)** Primary rat hippocampal neurons that had been cultured for five days were co-transfected with a plasmid encoding a fusion of the tail domain of KIF13A with green fluorescent protein (GFP) and a plasmid encoding Halo-tagged TfR. At 6 h after transfection, cultures were imaged for 30 sec at 2 frames/sec. The Halo tag was visualized using dye JF594. (A) Single frames of the time series showing the distribution of the tail domain of KIF13A and TfR. In the kymograph at the *bottom right,* the green line corresponds to the tail domain; the red line, to TfR. AIS, axon initial segment; Dend, dendrite. Scale bar, 25 μm. (B) Quantification of co-localization between the tail domain of KIF13A and TfR. **(C-E)** Primary rat hippocampal neurons that had been cultured for five days were co-transfected with a plasmid encoding guide RNA (gRNA1 or gRNA2) targeting KIF13A and a plasmid encoding a fusion of green fluorescent protein to the transferrin receptor (TfR). (C) Single frames of the time series showing the distribution of TfR in axons and dendrites. Kymographs are shown next to the high maleficiated images. Horizontal scale bar, 25 μm; vertical scale bar, 5 μm. (D) Quantification of vesicles containing TfR. (E) Quantification of the polarization of transport of vesicles containing TfR. ns, not significant; *P < 0.05, ****P < 0.0001 (Student’s *t* test).

To test if KIF13A transports vesicles containing TfR, small-guide RNA (KIF13A-gRNA) was designed in KIF13A exons. With the present of CRISPR-Cas9, the expression of KIF13A was suppressed (**Figure S1A)**. We then used KIF13A-gRNA to suppress KIF13A in neurons and recorded the trafficking of TfR vesicles. For a short period of time (1 day after transfection), we did not see an obvious change in the trafficking of TfR vesicles. After 3 days, we observed a decrease in the trafficking of TfR vesicles **(Fig1C, D)**. In the control neuron where scramble-gRNA were transfected, the trafficking of TfR vesicles were not disturbed. Although knockout of KIF13A largely impaired the transport of TfR vesicles, it did not change the polarization of of TfR vesicles, which probably means that dynein aggregated in the AIS region recycle TfR vesicles to the soma. Analysis was represented by the dendrite-axon ratio of TfR intensity (**Figure 1C, E**).

### Overexpression of KIF13A allows axonal transport of vesicles containing transferrin receptor

To test how KIF13A may affect the polarized transport of vesicles containing TfR, we overexpressed KIF13A in primary hippocampal neurons and recorded the movement of vesicles containing TfR. Overexpressed KIF13A leads to substantial traffic of vesicles containing transferrin receptor into axons, reducing the dendrite-axon ratio of TfR intensity (**Figure 2A-B, Movie 2**). As expected for the direction of KIF13A migration along microtubules, most vesicles in the axon moved anterograde (**Figure 2C**). The loss of purely dendritic polarity of vesicle transport was associated with reduced uptake of transferrin (**Figure S2A**), which may be a consequence of the mis-localization of transferrin receptor.

**Figure 2.**
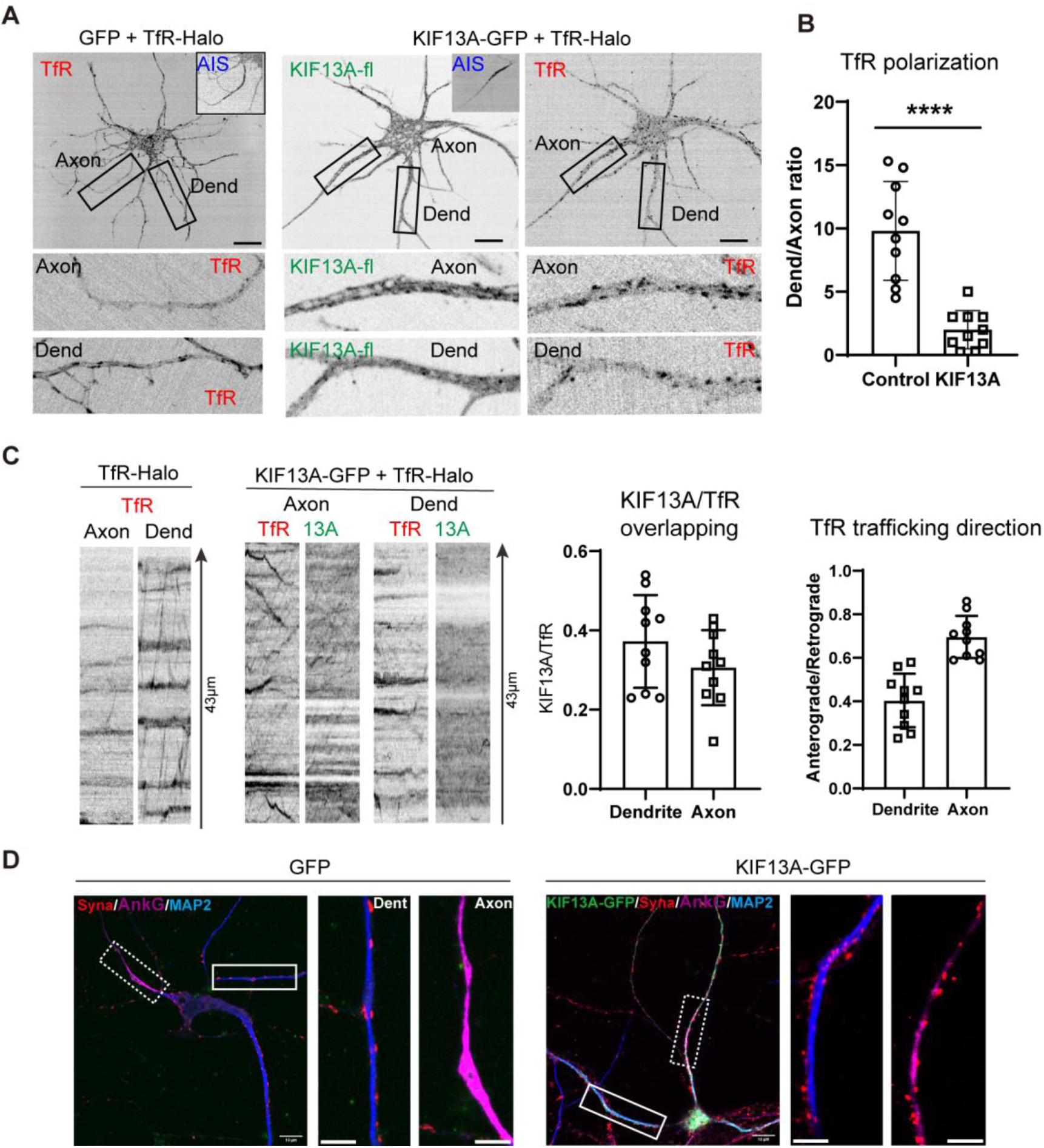
Overexpression of KIF13A in primary hippocampal neurons allows vesicles containing transferrin receptor (TfR) into axons. **(A-C)** Rat hippocampal neurons that had been cultured for five days were co-transfected with a plasmid encoding Halo-tagged transferrin receptor (TfR) and a plasmid encoding either a fusion of green fluorescent protein (GFP) with full-length KIF13A or GFP on its own. At 6 h after transfection, cultures were imaged for 30 sec at 2 frames/sec. (A) Single frames of the time series. Some frames contain insets showing the axon initial segment (AIS). Dend, dendrite. Scale bar, 20 μm. (B) Quantification of the polarization of transport of vesicles containing TfR. Control expresses GFP instead of KIF13A-GFP. (****P < 0.0001 based on Student’s *t* test.) (C) Kymographs of TfR-containing vesicles in dendrites or axons on the *left,* and quantification of co-localization between KIF13A and TfR as well as TfR trafficking direction on the *right.* (D) Rat hippocampal neurons that had been cultured for five days were co-transfected with a plasmid encoding either a fusion of GFP with full-length KIF13A or GFP on its own. After another nine days in culture, neurons were fixed and stained against synaptophysin (Syna) as a synaptic marker, ankyrin G (AnkG) as a marker of the axon initial segment, or MAP2 as a marker of dendrites. Scale bar, 10 μm (whole-cell image at *left)* or 5 μm in the zoomed views (at *right)*.

Moreover, we found that overexpression of KIF13A affected the polarity of endogenous synapses proteins (**Figure 2D**). We overexpressed KIF13A in cultured hippocampal neurons at DIV5 and stained synaptophysin at DIV14 for presynaptic particles. In control neuron transfected with GFP, synaptophysin staining was mainly at the dendrites (labeled with dendrite marker: Map2). In KIF13A-GFP transfected neurons, synaptophysin particles were observed in the axon (labeled with axon marker: ankyrin G), suggesting that the overexpressed KIF13A shifted the polarity of synapses protein from dendritic to partially axonal.

### Polarity of trafficking of vesicles containing transferrin receptor depends on KIF13A motor activity

Newly synthesized proteins are sorted at the Golgi complex into distinct vesicle populations. Then vesicles containing dendritic proteins or axonal proteins are transported to different area by kinesins (5). Disturbing the sorting process could result a miss targeting of dendritic protein to axonal vesicles. To test if overexpressed KIF13A miss-sorted TfR to axonal vesicles, we labeled axonal vesicles with neuron-glia cell adhesion molecule (NgCAM) and checked TfR distribution with or without KIF13A **(Figure 3A, Movie 3&4)**. Vesicles containing NgCAM showed a high preference to the axon and excluded from vesicles containing TfR. When KIF13A was overexpressed, vesicles containing TfR started moving into the axon, but no TfR contamination was observed in vesicles containing NgCAM in the axon. This indicates that overexpressed KIF13A changes the trafficking of vesicles containing TfR, while it does not affect the sorting of TfR.

**Figure 3.**
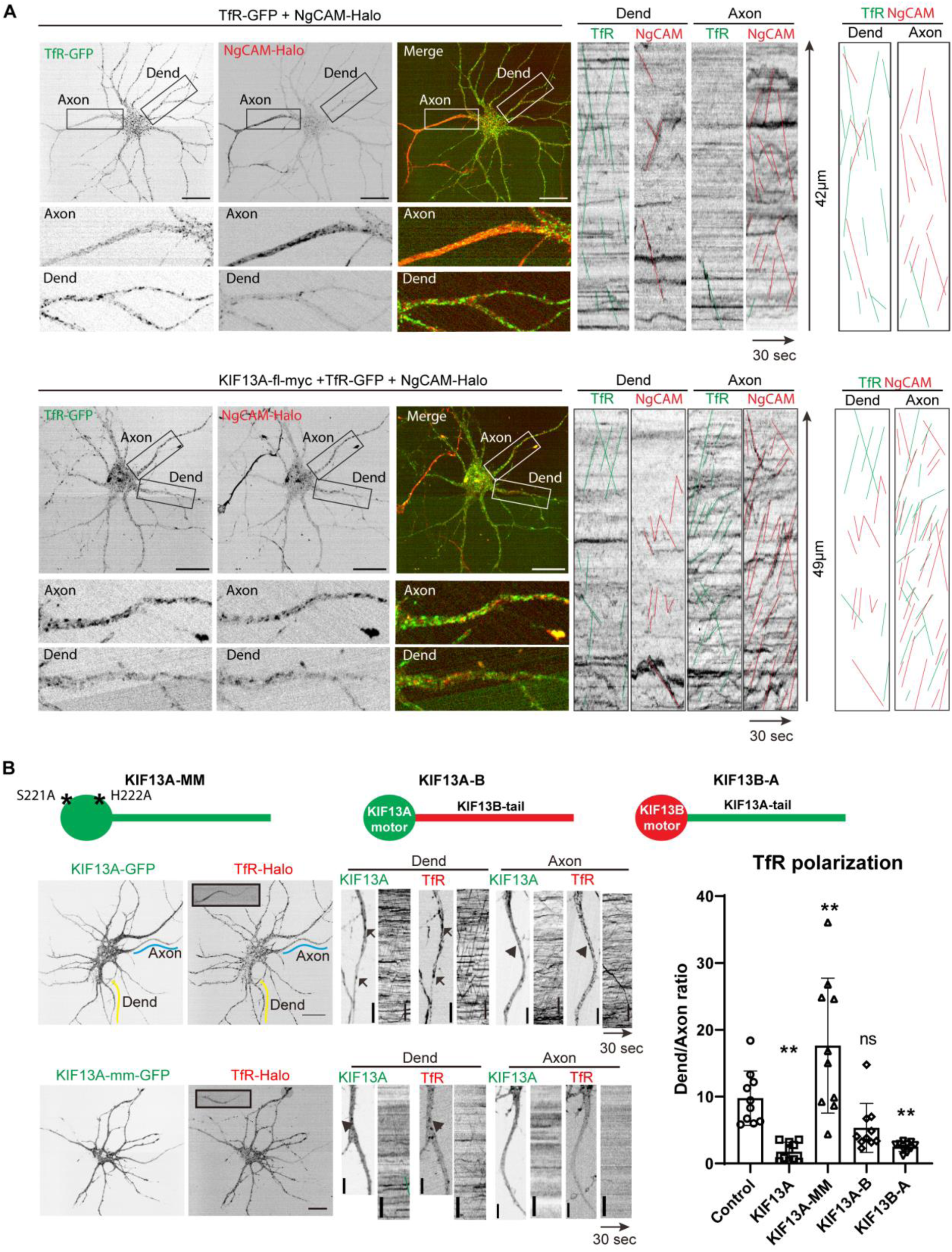
A tail domain of KIF13A is required to shuttle vesicles containing transferrin receptor (TfR) to axons. **(A)** Rat hippocampal neurons that had been cultured for five days were co-transfected with a plasmid encoding a fusion of green fluorescent protein (GFP) with TfR, a plasmid encoding Halo-tagged NgCAM and with or without a plasmid encoding myc-tagged full-length KIF13A. At 6 h after transfection, cultures were imaged for 30 sec at 2 frames/sec. The Halo tag was visualized using dye JF594. Single frames of the time series are displayed on the left. Kymographs of vesicles are displayed on the *right,* where green lines represent TfR vesicles and red lines represent NgCAM vesicles. Dend, dendrite. Scale bar, is 25 μm. **(B)** Rat hippocampal neurons that had been cultured for five days were co-transfected with a plasmid encoding Halo-tagged TfR and one of the indicated kinesin constructs: KIF13A-mm, containing two point mutations (Ser221Ala, His222Ala; labeled as asterisks) to abrogate ATPase activity of the motor domain; a chimera of the motor domain of KIF13A with the tail domain of KIF13B; or a chimera of the tail domain of KIF13A with the motor domain of KIF13B. As a control, some neurons expressed a fusion of GFP with wild-type, full-length KIF13A. At 6 h after transfection, cultures were imaged for 30 sec at 2 frames/sec. Typically micrographs are shown, and the staining of axon initial segment by antibody against neurofascin is displayed as an inset at the *upper left*. The extent of polarized trafficking of TfR vesicles is quantified on the *right*. Horizontal scale bar, 25 μm; vertical scale bar, 5 μm.

We next investigated whether motor activity is required for driving vesicles containing TfR to the axon. We introduced mutations (Ser221Ala, His222Ala) to the motor domain of KIF13A (KIF13A-mm) to unable to hydrolyze ATP (13, 14). Overexpression of KIF13A-mm did not induce axonal delivery of vesicles containing transferrin receptor (**Figure 3B**). In fact, it led to an even higher ratio of dendritic to axonal vesicle delivery than in wild-type primary hippocampal neurons.

Expression of a fusion of the motor domain of KIF13A with the tail domain of KIF13B, another kinesin-3 family member that may promote dendritic trafficking of vesicles other than those containing transferrin receptor (1), led to a ratio of dendritic to axonal delivery of transferrin receptor-containing vesicles similar to the ratio in wild-type neurons. In contrast, expression of a fusion of the motor domain of KIF13B with the tail domain of KIF13A allowed axonal vesicle transport similar to that in neurons overexpressing wild-type KIF13A (**Figure 3B**).

These results suggest that specifically KIF13A mediates the polarized delivery of vesicles containing transferrin receptor to dendrites, and that this depends on the motor activity of KIF13A.

### MARK2 phosphorylates KIF13A at a binding site for protein 14-3-3

Next, we explored what upstream proteins might regulate the effect of KIF13A on polarized vesicle transport. We engineered a plasmid encoding the tail domain of KIF13A fused to *E. coli* biotin ligase BioID2 (15), expressed the plasmid in HEK293T cells and screened the lysate for biotinylated proteins, which were candidate interactors of KIF13A (16). We detected numerous potential interactors (**Figure S4A-C, Table 1**), among which we focused on “microtubule affinity-regulating kinase 2” (MARK2), also known as Par1, because it has been implicated in neuronal polarity (17, 18). In addition, Par1b/MARK2 phosphorylates KIF13B at a binding site for protein 14-3-3 in order to regulate axonal formation (19).

First, we demonstrated that MARK2-GFP is enriched in the axon initial segment in primary hippocampal cultures (**Figure 4A**). The distribution of MARK2 in neuron is similar to neurofascin 186, the neuronal form of NF, which specifically localizes in the initial segment of axon and Ranvier node. Co-immunoprecipitation of MARK2 using subfragments of the tail domain of KIF13A identified the region encompassing residues 915-1749 in KIF13A as responsible for interaction with the kinase (**Figure 4B**). This was also the only subfragment of KIF13A that co-localized with vesicles containing transferrin receptor in primary hippocampal neurons (**Figure 4C-D**). A different subfragment of KIF13A, encompassing residues 362-914, also co-localized with vesicles in neurons, but not with the subset of vesicles containing transferrin receptor.

**Figure 4.**
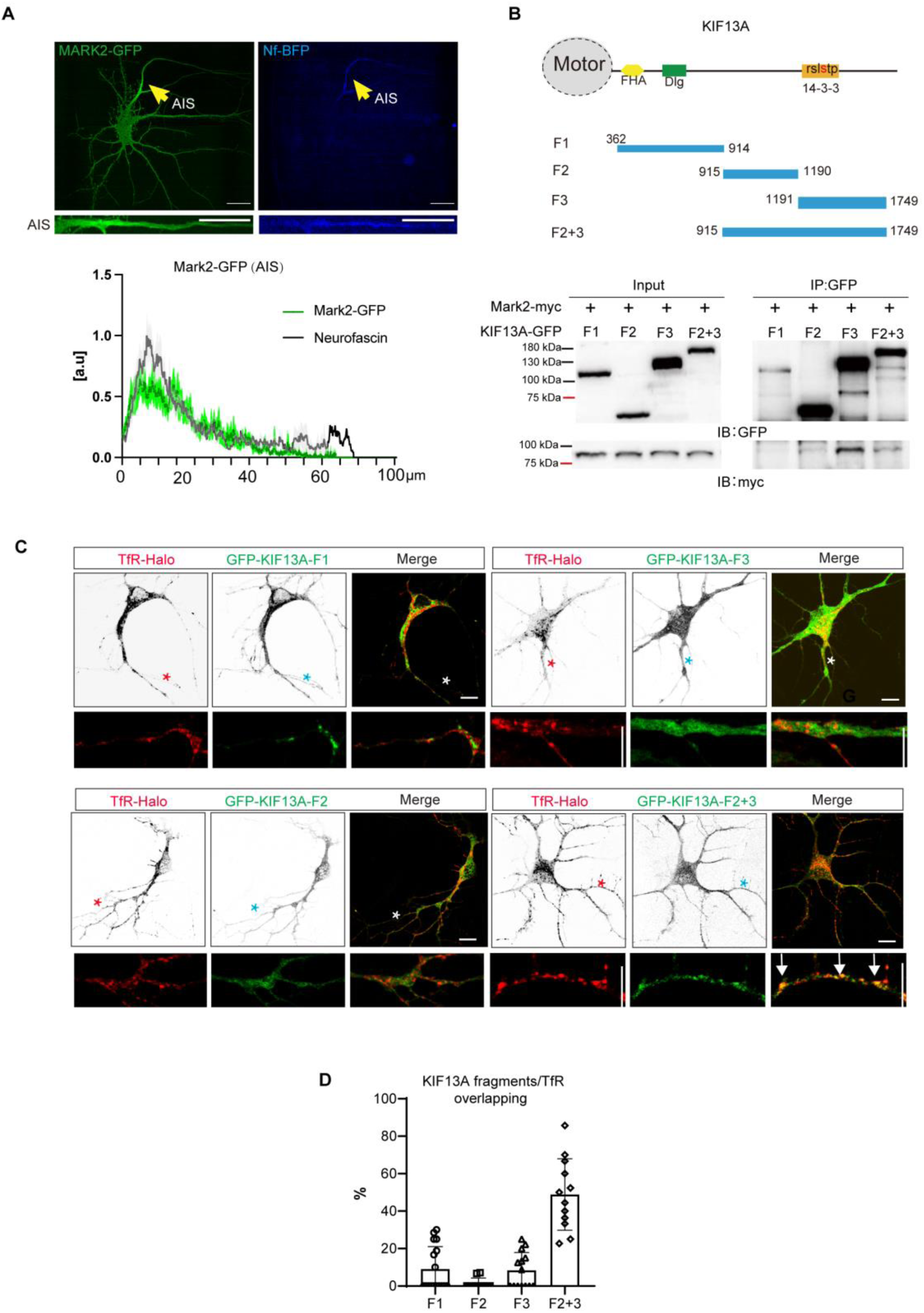
MARK2 binds to and phosphorylates the tail domain of KIF13A. **(A)** Rat hippocampal neurons that had been cultured for five days were co-transfected with a plasmid encoding a fusion of green fluorescent protein (GFP) with MARK2. The axon initial segment was stained with antibody against neurofascin conjugated with Alexa Fluor™ 405 (Nf-405). The relative localization of each protein relative to the soma is shown *below*. AIS, axon initial segment. Scale bar, 25 μm (*upper* micrographs) or 5 μm (*lower* micrographs). **(B)** The indicated fragments of the tail domain of GFP-tagged KIF13A were co-expressed with myc-tagged MARK2 in HEK293T cells, then the tail domain was immunoprecipitated from cell lysates using an anti-GFP antibody. The immunoprecipitates were probed using an anti-myc antibody. FHA, forkhead-associated; 14-3-3, binding site for protein 14-3-3. **(C-D)** Rat hippocampal neurons that had been cultured for five days were co-transfected with a plasmid encoding Halo-tagged transferrin receptor (TfR) and a plasmid encoding a fusion of GFP with one of the tail domain fragments in panel B. Asterisks mark dendrites shown at higher magnification in the *lower* images, where white arrows indicate co-localization of KIF13A tail and TfR. Horizontal scale bar, 25 μm; vertical scale bar, 5 μm. (D) Quantification of co-localization.

Given that MARK2 has been shown to phosphorylate the tail domain of KIF13B at a binding site for the protein 14-3-3 (_1377_RXXXSXP_1383_) (19), we wondered whether the same might be true of KIF13A. Indeed, we identified a conserved binding site for protein 14-3-3 in the tail domain, (_1367_RSLSTP_1373_) (**Figure 5A**), which coincided with the fragment that we identified above as critical for interaction with MARK2. Then we mutated the Ser1371 at the phosphorylation site to Asp in order to mimic phosphorylation, or to Ala to prevent phosphorylation. In HEK293T cells, the Ser1371Asp mutant of KIF13A bound to 14-3-3ε much more strongly than the Ser1371Ala mutant did, as well as isoform ζ or isoforms β (**Figure 5C**). These results suggest that the phosphorylation of KIF13A at the 14-3-3 binding site by MARK2 strengthens the kinesin’s interaction with 14-3-3.

**Figure 5.**
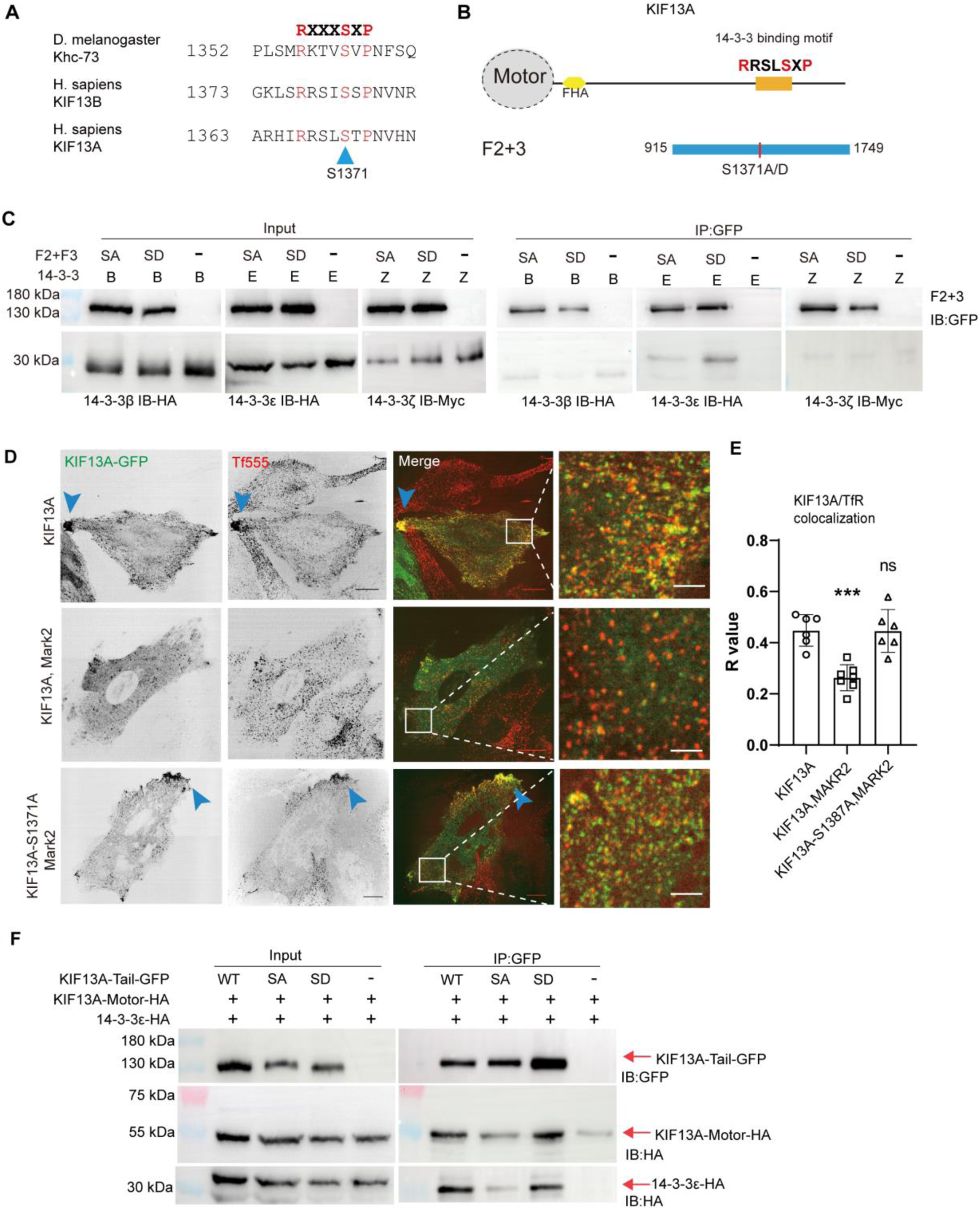
Phosphorylation of Ser1371 in the tail domain of KIF13A by MARK2 is required for the kinesin to associate with vesicles containing transferrin receptor (TfR). **(A)** Conservation of the 14-3-3 binding motif across species. The consensus sequence is written at the *top*. The Ser1371 in KIF13A that MARK2 phosphorylates is indicated at the *bottom*. **(B)** Construction of a fragment of the tail domain of KIF13A that combines fragments F2 and F3 in Figure 4B and contains either a mutation to remove the MARK2 phosphorylation site (Ser1371Ala) or a mutation to mimic phosphorylation (Ser1371Asp). **(C)** HEK293T cells were co-transfected with a plasmid expressing a fusion of green fluorescent protein (GFP) with Ser1371Ala or Ser1371Asp mutants of the tail domain of KIF13A; and with a plasmid expressing myc-tagged 14-3-3 isoform ζ or HA-tagged isoforms ε or β. KIF13A fragments were immunoprecipitated using anti-GFP antibody, and the immunoprecipitates were probed with anti-myc or -HA antibody. **(D-E)** Rat embryonic fibroblasts were transfected with a plasmid expressing wild-type KIF13A full-length or the Ser1371Ala mutant and, in in some cases, with a plasmid expressing MARK2. Cultures were treated with transferrin conjugated to the fluorescent dye Alexa 555 (Tf555). (D) Representative fluorescence images showing the localization of KIF13A and TfR vesicles. The area boxed in white is shown at higher magnification at the *far right*. Scale bar, 25 μm or 5 μm (*far right*). (E) Co-localization of KIF13A and TfR vesicles. (***P < 0.001 based on Student’s *t* test.) **(F)** HEK293T cells were co-transfected with a plasmid encoding a fusion of GFP with the wild-type or mutant version of the tail domain of KIF13A, along with a plasmid encoding the HA-tagged motor domain of KIF13A and the HA-tagged 14-3-3 isoform ε. KIF13A fragments were immunoprecipitated using anti-GFP antibody, and immunoprecipitates were probed using anti-HA antibody. ns, not significant; ***P < 0.001 (based on Student’s *t* test).

### Phosphorylation of KIF13A by MARK2 detaches the kinesin from vesicles containing transferrin receptor

We examined the association of KIF13A with vesicles containing transferrin receptor and the subsequent trafficking of those vesicles in rat embryonic fibroblasts. These cells are flatter than neurons, which facilitates detection of kinesin co-localization with vesicles; and the microtubules in these fibroblasts radiate uniformly outward from the centrosome to the cell periphery, which facilitates observation of the direction of kinesin movement. Vesicles containing transferrin receptor in these fibroblasts were evenly distributed throughout the cytosol in the absence of KIF13A, while co-expression of the kinesin caused the vesicles to accumulate at the cell periphery (**Figure 5D**). This accumulation at the cell periphery was partially reversed by co-expressing MARK2 with KIF13A, which also reduced co-localization between KIF13A and vesicles (**Figure 5E**). These effects of MARK2 were abrogated when KIF13A contained the Ser1371Ala mutation preventing phosphorylation. These results suggest that phosphorylation of KIF13A by MARK2 inhibits the kinesin’s ability to interact with, and traffic, vesicles containing transferrin receptor.

In support of this idea, we found that in HEK293T cells expressing separate motor and tail domains of KIF13A, the motor domain of KIF13A interacted more strongly with tail domains containing the wild-type 14-3-3 binding motif or the motif with the Ser1371Asp mutation mimicking phosphorylation than with a tail domain containing the non-phosphorylatable Ser1371Ala mutation (**Figure 5F**). Since stronger interaction between the motor and tail domains of kinesin inhibits its binding to microtubules (20), our findings argue that phosphorylation of KIF13A by MARK2 detaches the kinesin from vesicles containing transferrin receptor.

### Phosphorylation of KIF13A by MARK2 is required for dendritic trafficking of vesicles containing transferrin receptor

To clarify how phosphorylation of KIF13A by MARK2 affects polarized transport of vesicles containing transferrin receptor, we knocked out MARK2 using CRISPR-Cas9 (**Figure S4**) and analyzed the trafficking of such vesicles in primary cultures of hippocampal neurons (**Figure 6A-B**). Knockout increased the proportion of vesicles trafficked into axons rather than dendrites, consistent with what we observed in rat embryonic fibroblasts expressing the Ser1371Ala mutant of KIF13A. Indeed, again consistent with our results in fibroblasts, we found that overexpressing KIF13A in primary hippocampal neurons expressing endogenous MARK2 allowed a certain degree of axonal trafficking of vesicles containing transferrin receptor (**Figure 6C-D**). The strictly dendritic trafficking of these vesicles was restored when KIF13A and MARK2 were overexpressed together, but not when the overexpressed MARK2 contained the active-site mutation Ser212Asp (MARK2 kinase dead, MARK2-KD) to abrogate its kinase activity or when the overexpressed KIF13A contained the non-phosphorylatable mutation Ser1371Ala.

**Figure 6.**
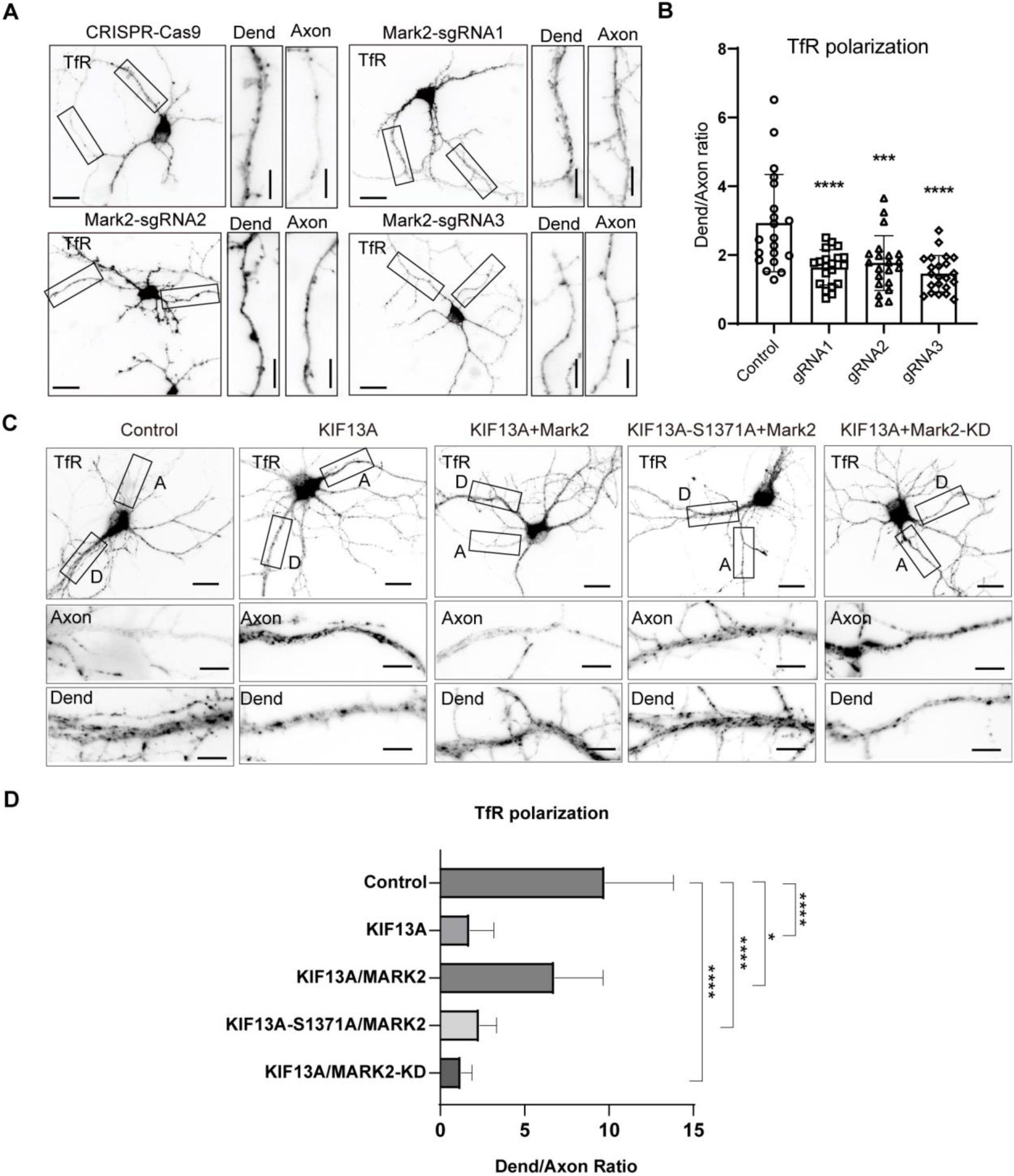
Phosphorylation of KIF13A by MARK2 is required for polarized transport of vesicles containing transferrin receptor (TfR) to dendrites. **(A-B)** Cultured hippocampal neurons at DIV5 were co-transfected with a plasmid expressing Halo-tagged TfR and a plasmid encoding the indicated small guide RNA (gRNA) targeting MARK2. (A) Representative photomicrographs. The areas boxed in black on the *left* are shown at higher magnification on the *right*. Dend, dendrite. Horizontal scale bar, 25 μm; vertical scale bar, 10 μm. (B) Extent of polarization of trafficking of TfR vesicles. **(C-D)** Cultured hippocampal neurons at five days were co-transfected with a plasmid expressing Halo-tagged TfR and a plasmid encoding a fusion of green fluorescent protein (GFP) with full-length wild-type KIF13A or the Ser1371A mutant. Some cultures were also transfected with a plasmid expressing wild-type MARK2 or the MARK2-Ser212Arg kinase-deficient (KD) mutant. The regions boxed in black *above* are shown at higher magnification *below*. A, axon; D or Dend, dendrite. Scale bar, 25 μm (*above*) or 10 μm (*below*). (D) Extent of polarization of trafficking of TfR vesicles. *P < 0.05, ***P < 0.001, ****P < 0.0001 (based on Student’s *t* test).

These results suggest that phosphorylation of KIF13A by MARK2 is required for the strictly dendritic trafficking of vesicles containing transferrin receptor in neurons.

## Discussion

Dendritic polarized vesicles were barely across the AIS and enter the axon. How neurons maintain dendritic vesicles is a key question in the neuron polarity field. Here we provide evidence that vesicle subpopulations containing transferrin receptor trafficked by the kinesin KIF13A are excluded from axons of neurons because the kinesin is phosphorylated by MARK2, which promotes its binding to 14-3-3 rather than to vesicles and inhibits the kinesin’s motor activity (**Figure 7**). Our results add KIF13A to the list of motor proteins that mediate polarized vesicle transport within neurons, a list that already includes myosin V (21) and dynein (22, 23)(24), as well as the dense network of actin filaments within the axon initial segment (25).

**Figure 7.**
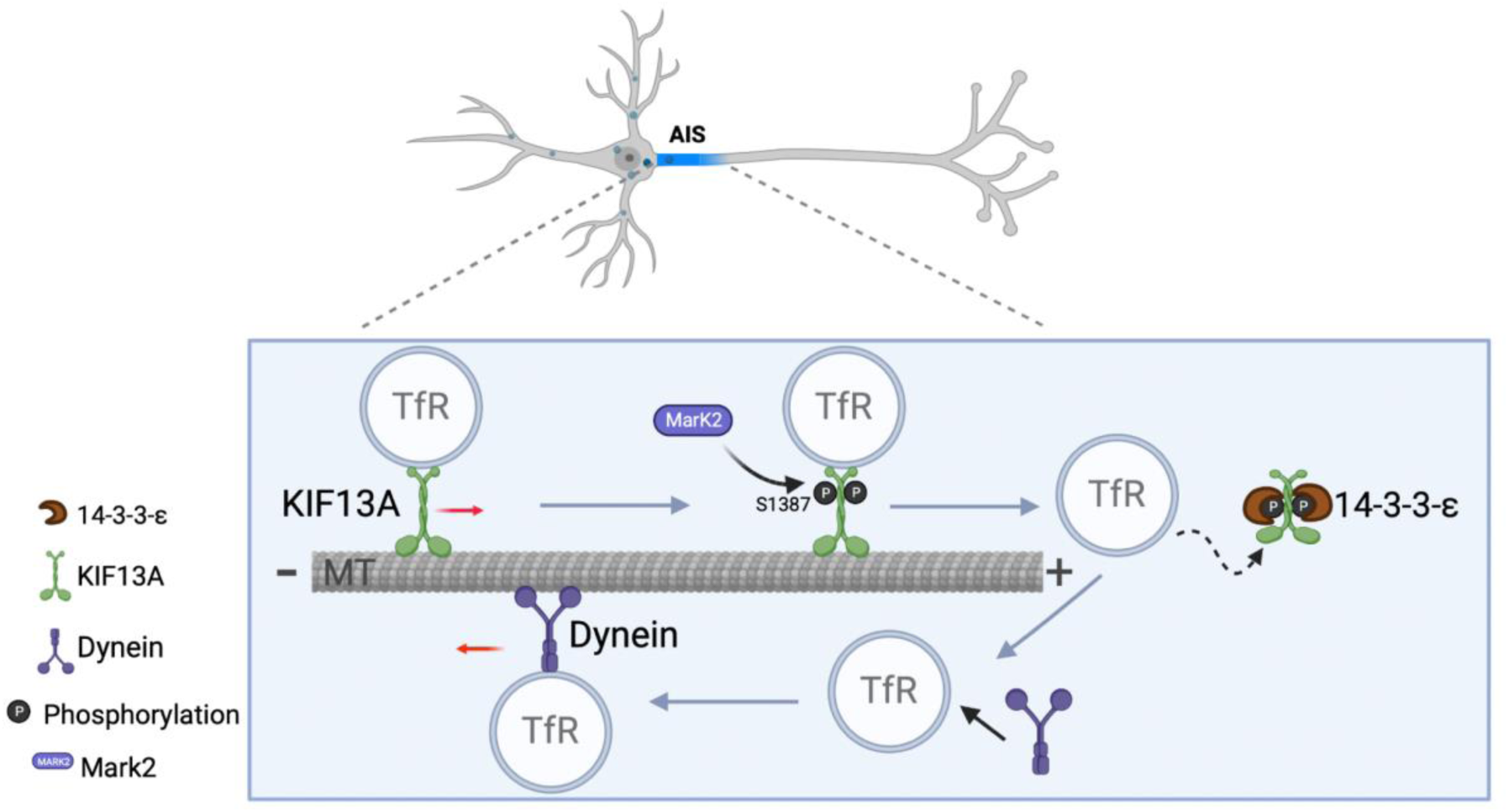
Model for phosphorylation of the 14-3-3 binding motif in KIF13A by MARK2 in the axon initial segment (AIS).

Since KIF13A shares high sequence homology with KIF13B, which also associates with vesicles containing transferrin receptor in neurons (26), we expected that the two kinesins would perform the same in our experiments. However, overexpressing fusion of the motor domain of KIF13A with the tail domain of KIF13B affected the polarization of such vesicles to a much weaker extent than the corresponding changes in fusion of the motor domain of KIF13B with the tail domain of KIF13A, suggesting that KIF13A is the dominant transporter of this vesicle subpopulation. Indeed, our previous work with primary hippocampal neurons showed that few vesicles transported by KIF13B contained transferrin receptor and that an appreciable proportion of KIF13B-associated vesicles travelled to the axon, whereas most vesicles transported by KIF13A contained transferrin receptor and few travelled to the axon (1). We hypothesize that the different tail domains of kinesin-3 family members confer selectivity for different subpopulations of vesicles, which should be explored in future work.

Previous studies have shown that Mark2 regulates neuronal polarization, especially the phosphorylation of KIF13B, with which KIF13A shares a conserved sequence. The interaction between Mark2 and KIF13A tail was confirmed by biotin affinity purification analysis. We found that overexpressing MARK2 in primary hippocampal neurons that had been cultured for one week led to the protein’s accumulation in the axon initial segment. In contrast, work of Hirokawa N and colleagues showed overexpressing MARK2 in primary neurons that had been cultured for four days led to its accumulation in soma and dendrites (27). This difference may reflect neurons at different developmental stages: the axon initial segment forms in primary neurons between four and seven days in culture (28). Future work should explore how MARK2 regulates polarized vesicle traffic at different stages of neuronal development.

14-3-3 is a conserved family of adaptor proteins that interact with diverse proteins and regulate their function. 14-3-3 family members were found interact with both dynein (29–31) and kinesin (32, 33). The interaction between 14-3-3 and dynein/kinesin could promote the dynein activity (29, 30) or inhibit kinesins activity. In this study, we showed phosphorylation of KIF13A by MARK2 enhanced the interaction between KIF13A and 14-3-3, which inhibited the kinesin activity. In the soma-dendritic area, due to the low abundance of MARK2 kinase, KIF13A bound to and actively transported TfR vesicles. At AIS, the highly expressed MARK2 phosphorylated and inhibited KIF13A, which paused TfR vesicles. In the meantime, dynein was activated and moved the vesicles back to the soma. Although how dynein is activated at AIS has been reported (22, 23), whether MARK2 is also involved in the regulation of dynein activity is a question worthy of follow-up study.

## Material and methods

### Primary cultures of hippocampal neurons

Procedures involving animals were approved by Xi’an Jiaotong university, basic medical school (Approval number:2022-0046). Astroglia cells and pyramidal neurons were isolated from 1-day-old Sprague-Dawley rat pups as described (34). In brief, the rat brain was placed in a dish containing CMF-HBSS (Gibco, Catalog number: 14175095), meninges were stripped away, then the cortex was isolated and chopped up finely and transferred to a 50-ml centrifuge tube containing 15 ml lysis buffer [12.5 ml CMF-HBSS, 1.5 ml 2.5% trypsin, 1 ml 1% (w/v) DNase]. The tissue was lysed by incubating in a water bath at 37 ℃ for 5 min (with swirling every 2 min), pipetting up and down in a 10-ml pipette 10-15 times, incubating at 37 ℃ another 10 min (with swirling every 2 min), then pipetting up and down with a 5-ml pipette until obvious chunks had dispersed. The lysate was passed through a cell strainer, collected into a 50-ml tube containing equal volumes of glial medium (DMEM supplemented with Penicillin-Streptomycin and 10% horse serum), and centrifuged for 5 min at 120 *g.* The cell pellet was resuspended with glial medium, seeded into 9-cm wells (9X10^6^ cells per well), and incubated 2 weeks to serve as a feeder layer for primary hippocampal neurons. Culture medium was changed on the next day every 3 days following seeding.

To extract hippocampal neurons, we harvested hippocampi from 1-day-old rat brains into a 15-ml tube containing approximately 10 ml of CMF-HBSS. Hippocampi were rinsed twice with CMF-HBSS, transferred to a fresh 15-ml tube containing lysis buffer [4.5 ml CMF-HBSS, 0.5 ml 2.5% trypsin], incubated in a water bath at 37 ℃ for 15 min, and remove the solution. The pellet was resuspended in 5 ml CMF-HBSS and allowed to stand at room temperature for 5 min. This procedure was repeated once, then the pellet was resuspended in only 1-2 ml of solution and passed 10 times through a Pasteur pipette with a standard diameter, followed by 10-20 times through a Pasteur pipette with 50% smaller diameter. Adjust the cell density to about 50,000 to 500,000 cells per prepared diameter 6-cm dish contain 5-6ml plating medium [DMEM supplemented with 0.6% glucose and 10% horse serum]. The cell dish containing 3-4 glass coverslips (which were cleaned and coated with poly-L-lysine before 2 days). After 2–4 h, transfer the coverslips to the glial feeder layer described above. The glial feeder dish contained 10-12ml Neurobasal medium (Gibco, Catalog number: 21103049) with B27 Supplement (Gibco, Catalog number: 17504-044) and GlutaMAX-1 Supplement (Gibco, Catalog number: 35050079).

### Knockout of KIF13A or MARK2 in primary hippocampal neurons

The CRISPR-Cas9 system was used to knockout KIF13A or MARK2 in primary hippocampal neurons using a short guide RNA targeting exons 5, 6 in KIF13A, and exons 2, 3, 6 in MARK2. The short guide RNA was cloned into the vector lentiCRISPRv2 (35, 36). Primary neurons were cultured for 48 h as described above, then co-transfected for 2 days with the lentiviral plasmid and plasmid encoding a fusion of Halo to the transferrin receptor (The plasmid was from Dr. Gary Banker’s lab in OSHU) (0.25μg per coverslip for each plasmid). Knockout effect of KIF13A or MARK2 were confirmed by western blot. In brief, lentiCRISPRv2-gRNA plasmids of KIF13A or MARK2 were co-transfected with KIF13A-fl-myc or MARK2-GFP into 293T cells for 48h, then harvest the cells with 2X SDS loading buffer, then detecting Myc and GFP labeling of KIF13A and MARK2.

### Construction of plasmids to express domains and mutants of KIF13A or MARK2 in primary neurons or HEK283T cells

Plasmids expressing the tail domain of KIF13A were generated by replacing the N-terminal motor domain with enhanced green fluorescent protein, while plasmids expressing the motor domain of KIF13A were generated by replacing the C-terminal tail with a hemagglutinin (HA) tag. In construction of mutant tail of KIF13A, briefly, the serine of 1371site is replaced with either suppressed arginine or activated aspartic acid. Constructs are described in **Supplementary Table 1**.

### Transfection of primary neurons and HEK283T cells

Primary neurons were transfected with expression plasmids using Lipofectamine 2000 (Invitrogen). In brief, primary neurons were removed and placed on a new dish which contained 70% fresh neurobasal medium with B27 and GlutaMAX-1 and 30% neurobasal medium from original glial feeder dish before transfection. Expression plasmids including KIF13A-fl-GFP, TfR-GFP, NgCAM-Halo, KIF13A-fl-myc, KIF13A-MM-GFP, KIF13A-B-GFP, KIF13B-A-GFP, Mark2-GFP, KIF13A-S1371A, pEGFP-KIF13A Tail (F1, F2, F3, F2+3) and TfR-Halo were mixed with Lipofectamine 2000 (0.25μg per coverslip for each plasmid) in Opti-MEM (Gibco, Catalog number: 31985070), transfer the mixture into primary neurons for 30 min, then seeded onto the glial feeder layer described above. HEK283T cells were transfected using calcium phosphate precipitation as follows. When cells were 40-60% confluent, they were fed for fresh growth medium with DMEM containing 5% fetal bovine serum, L-glutamine and penicillin/streptomycin. pEGFP-KIF13A Tail/Mark2-myc, pEGFP-KIF13A Tail/KIF13A Motor-HA, pEGFP-KIF13A Tail/14-3-3ε-HA, pEGFP-KIF13A Tail/14-3-3ζ-myc and pEGFP-KIF13A Tail/14-3-3β-HA expression plasmids (5 μg per 10-cm plate for each plasmid) were suspended in 12.4% 2 M CaCl_2_, which was added dropwise to isopycnic 2x HBS (Containing 10mM D-Glucose,40mM HEPES, 10mM KCl, 270mM NaCl and 1.5mM Na_2_HPO_4_) during fewer than 5 min. The resulting solution was added to the cells, the plates were shaken gently, then the cultures were incubated 12-16 h. Fresh growth medium was added, then cultures were incubated in incubator till use.

### Immunoprecipitation of Mark2/KIF13A motor/14-3-3 in HEK293T cells

HEK293T cells that had been transfected with expression plasmids as described above were centrifuged at 120 g for 5 min at 4 °C, and the pellet was resuspended in 500 μl ice-cold lysis buffer (Containing 500mM NaCl, 0.5mM EDTA,10mM HEPES, 0.5% NP-40, 50mM Sodium Fluoride 10mM Sodium Pyrophosphate and Pierce™ Proteinase inhibitor, #A32963; Thermo Scientific, TX, USA). Cells were passed through a syringe (0.6X25mm) 5-10 times, incubated on ice for 30 min, and centrifuged at 14,000 *g* for 30 min at 4 °C. The supernatant was transferred to a fresh, pre-chilled tube. An aliquot of supernatant (50 μl) was mixed with 5X loading buffer to serve as the “input” sample. The remaining supernatant was subjected to immunoprecipitation using 15 μl of mouse monoclonal antibody against green fluorescent protein (catalog no. sc-9996, Santa Cruz, CA, USA). The immunoprecipitate was gently shaken overnight at 4 °C, then mixed with 15 of μL magnetic bead slurry (Pierce™ Protein A/G Magnetic Beads, catalog no. 88802; Invitrogen, CA, USA) in a pre-chilled tube, incubate for 1-2 hours at 4°C with gentle agitation. Then, the suspension was washed with 500 μL of wash buffer (Containing 150mM NaCl, 0.5mM EDTA,10mM HEPES, 0.05% NP-40), pipetted up and down several times, then the beads were pulled down with a magnet. This procedure was repeated twice more. Then the beads were mixed with 50 μL of cell lysis buffer, and the supernatant was further analyzed by immunoblotting.

### Time-lapse imaging

We imaged primary cultures as described (1). Cells were incubated while imaging in Hibernate E medium without phenol red (Brain-Bits), then images were acquired with an sCMOS camera (Zyla 4.2+, Andor) under a Ti-E microscope (Nikon) equipped with a spinning-disk confocal head (model CSU-W1, Yokogawa). Vesicles containing transferrin receptor were imaged using an sCMOS camera through a CFI Apo 60× 1.49 objective (Nikon) on a Ti2 microscope (Nikon). The entire imaging stage and objectives were maintained at 37 °C in a closed system (customized by OkoLab). Image streams (two frames per sec) were acquired using a Plan-Apo 100× 1.49 NA objective (Nikon) with 2 × 2 binning, during which z-axis movement was controlled using the Perfect Focus system on the Ti-E microscope (Nikon).

### Immunocytochemistry of primary hippocampal neurons cells

Cells on coverslips were washed with PBS, incubated at 37 ℃ for 10 min in pre-warmed fixation buffer containing 4% (w/v) paraformaldehyde + 4% (w/v) sucrose in 0.01 M PBS, washed three times with PBS, permeabilized with 0.25% Triton X-100 for 10 min, washed again three times with PBS, incubated with desired antibody at 4 degree, overnight incubation. For conditions where Halo tag was used, cells were incubated with 20 nM JF-549® dye in 1xPBS for 10 min at room temperature. After washed with PBS, cells were incubated with desired secondary fluorescent antibody for 1 hour at room temperature. Washed with 1xPBS 3 times and fixed with mounting medium (ProLong™ Diamond; Thermo Fisher Scientific, P36961) for imaging. Imaging was taken with fv3000 olympus microscopy. The antibodies used for immunostaining are described in **Supplementary Table 2.**

### Statistical analysis

Kymographs were generated from image series using Fiji 2.12, and the numbers and velocities of puncta were compared between axons and dendrites using Student’s *t* test. Only puncta that could be unambiguously assigned as vesicles were included in the analysis. Functional enrichment analysis of the biotinylated proteins was performed using RStudio version 2023.06.2+561. Missing values of area in list obtained after pre-quality control screening (Table 1) were imputed as the mean value of group. In order to ensure the comparability of proteins, the KIF13A area of each group were normalized, and the area of each protein was standardized according to this standard. Significantly enriched proteins (GFP-KIF13A Tail-BioID *vs* GFP-BioID, T test unadjusted p≤0.05) were analyzed by the Go_enrich program. In assays involving Primary pyramidal neurons in which MARK2 was knocked out, polarization between axons and dendrites was semi-quantified in terms of the ratio of fluorescence in axons to the fluorescence in dendrites.

## Acknowledgments

This work was initiated at the Jungers Center for Neurosciences Research, Oregon Health & Science University, Portland, Oregon when I was a postdoc under Prof. Gary Banker’s supervision. We thank Prof. Marvin Bentley for his generous help in experiment design and interpretation of the results. We thank Julie Luisi and Zoe Bostick for plasmid construction and primary culture of hippocampal neurons. We thank Prof. Wei Feng at National Laboratory of Biomacromolecules Institute of Biophysics, Chinese Academy of Sciences for providing us the KIF13A plasmid. We thank Prof. Yongqi Huang at Key Laboratory of Industrial Fermentation, Hubei University of Technology for providing us the 14-3-3 plasmids. We thank Dr. Qiaoyi Chen for grammar corrections on the manuscript. The portion of this work performed at Oregon Health and Sciences University was supported by National Institutes of Health grant MH066179 to Marvin Bentley. The work performed at Xi’an Jiaotong University was supported by the National Natural Science Foundation of China (33271016 to RY, 82100454 to JXX, 82101586 to HY).

